# *De novo* genome sequence assembly of the model algal endosymbiont *Micractinium conductrix* derived from its host *Paramecium bursaria* 186b

**DOI:** 10.64898/2025.12.15.694297

**Authors:** Guy Leonard, Irma Vitonytė, Fiona R. Savory, Erika M. Hansson, Duncan D. Cameron, Michael A. Brockhurst, Thomas A. Richards

## Abstract

Endosymbiosis is a major driver of evolutionary innovation and underpins the function of diverse ecosystems. The origins and evolution of endosymbiosis are challenging to study experimentally due to the short-lived culturability of many microbial strains derived from endosymbiotic interactions. The facultative endosymbiosis between the ciliate, *Paramecium bursaria*, and the green alga, *Micractinium conductrix* (Chlorellaceae, Trebouxiophyceae), is ecologically widespread and has emerged as a powerful lab-tractable model system. This endosymbiosis is founded upon a reciprocal nutrient exchange, but each of the species can be cultured independently enabling quantification of symbiotic fitness effects, new partnerships to be generated in the lab, and co-associations to be subject to experimental evolution. To date, evolve-and-resequence approaches have been limited due to a lack of high-quality genome assemblies enabling gene variants to be identified. Here, we report a near telomere-to-telomere genome assembly for *M*. *conductrix* 186b, using a range of sequencing technologies. Comparative analysis shows that this is one of the most complete Chlorellaceae algal genome assemblies available to date. To aid accurate gene calling and annotation we conducted both RNAseq and Iso-Seq transcriptome sequencing experiments. Collectively these ‘omics datasets will facilitate: i) comparative genomics studies of endosymbiont evolution, ii) evolve-and-resequence experiments, iii) genome-scale metabolic modelling studies, and iv) identification of targets for genetic modification experiments and biotechnological applications.

**Significance statement:** Endosymbiosis, where one species, the endosymbiont, lives inside the cell of another species, the host, has played a key role in evolution of complex life. However, the origins and evolution of obligate endosymbioses are often challenging to study because the key events are hidden deep in evolutionary time. Facultative microbial endosymbioses, such as between the ciliate *Paramecium bursaria* and the green alga *Micractinium conductrix*, offer experimentally tractable model systems where interacting species can be grown independently or in association allowing studies of the origin, evolution and fitness effects of symbiosis. Here we report the genome sequence for *M*. *conductrix* from *P. bursaria* 186b, enabling genomic studies of the evolution of endosymbiosis and simplifying gene target acquisition for microbe engineering and biotechnological applications.

## Introduction

Endosymbiosis is an important driver of evolutionary innovation and organismal complexity, enabling organisms to acquire novel functions and occupy new ecological niches (Szathmáry and Smith 1995; Douglas 2010; Wernegreen 2012; Brockhurst, et al. 2024). As such, endosymbiotic species interactions underpin ecosystem function across a wide range of biomes, including coral reefs and rain forests, as well as boosting the productivity of agricultural crops (Johnson 2015). Despite this wide-ranging importance, our understanding of the origins and evolution of the endosymbiotic process is limited because the key events are lost deep in evolutionary time (Brockhurst, et al. 2024). Moreover, for many extant endosymbiotic interactions, it is often highly challenging to culture the species independently, limiting direct experimental study. Facultative endosymbioses, where the interacting species retain the ability to grow independently, offer powerful experimentally-tractable tools for studying the evolution of endosymbiosis where fitness effects of symbiosis can be quantified. One such facultative endosymbiosis emerging as a key eco-evolutionary model is the nutritional endosymbiosis between the ciliate, *Paramecium bursaria* 186b, and the green algal, *Micractinium conductrix* 186b (Lowe, et al. 2016; Sørensen, et al. 2020; Jenkins, et al. 2021a; Jenkins, et al. 2021b; Sørensen, et al. 2021). A single *P*. *bursaria* cell contains several hundred algal cells arrayed around the interior surface of the host cell membrane, each individually packaged within a peri-algal vacuole, enabling the host to control nutrient and ion exchange (Jenkins 2024). This interaction is based upon a core nutrient exchange of algal photosynthate (sugars) for amino acids that hosts derive from heterotrophy (Sørensen, et al. 2020; Fujishima and Kodama 2022). Other potential benefits of this interaction include provisioning of oxygen from algal photosynthesis to the host, and protection from predation and viral infection of the alga by the host (Fujishima and Kodama 2012; Jenkins 2024).

Work by us, and others, has established a range of experimental methods that enable the detailed study of the cell biology, ecology, and evolution of this endosymbiotic interaction (Fujishima and Kodama 2012; Lowe, et al. 2016; Sørensen, et al. 2020; Jenkins, et al. 2021a; Jenkins, et al. 2021b; Sørensen, et al. 2021; Fujishima and Kodama 2022). For example, we have shown using lab ecological experiments that hosts tightly modulate symbiont load (i.e., the number of algal cells per host) in response to light intensity in order to maximise the benefit-to-cost ratio of the interaction (Dean, et al. 2016; Lowe, et al. 2016). Specifically, hosts reduce symbiont load both in the dark, where endosymbionts are purely costly, and at high light intensity, where fewer algal cells are required to supply the host’s carbohydrate requirement, minimising the cost to hosts of supplying endosymbionts with amino acids. Using partner-switching experiments we have shown that divergent host-symbiont pairings exchange a common core set of nutrients but have diverged in other traits linked to light management, such as photo-protectants and antioxidants (Sørensen, et al. 2020). Consequently, we observe specificity between hosts and algal symbionts, such that novel pairings of genotypes often have mismatched phenotype combinations, resulting in low fitness (Sørensen, et al. 2020). The genomic causes of this variation remain unknown, in part due to a lack of high-quality reference genomes for the algal endosymbiont. Using lab experimental evolution, we have shown that novel host-symbiont pairings with low fitness can rapidly attain high fitness through compensatory evolution, either via alterations to algal photosynthesis or host control of algal symbionts (Sørensen, et al. 2021). Re-sequencing of evolved host-symbiont genotypes would be required to understand the genetic adaptations that underly this rapid evolution, but is limited by a lack of reference genomes.

Many existing genome assemblies of *M. conductrix* and closely related algal endosymbionts are incomplete and/or highly fragmented (see Table 1). Here we present a replete ‘omics sequencing dataset and a high-quality assembled genome sequence, along with multiple transcriptome datasets, of the green algal endosymbiont *M. conductrix* 186b derived from its host *P. bursaria* 186b. We recently published the genome sequences of the host (Leonard, et al. 2025); this work therefore completes genome sampling of both partners involved in this endosymbiotic interaction. Using the data generated, we compare the new *M. conductrix* 186b genome assembly to other published Chlorellaceae genomes to understand differentially predicted gene fusions, a potential source of artefact in gene calling within this algal group.

**Table 1.** Comparison of genome assembly data. Data generated by reanalysis of all genomes using QUAST (Mikheenko, et al. 2018) and GAQET ((Mikheenko, et al. 2018) -images available in Fig. S2).

|  | *M. conductrix*<br>186b | *M. conductrix* SAG<br>241.80 | *M. sp. NFX-FRZ* | *M. tetrahymenae* | *M. sp.* CCAP<br>211/92 | *C. variabilis*<br>NC64A | *C. vulgaris* | *C. sorokiniana*<br>1602 | *C. sp* A99 |
| --- | --- | --- | --- | --- | --- | --- | --- | --- | --- |
| Citation | This paper | (Arriola, et al. 2018) | (Quintas-Nunes, et al. 2023) | (Kelly, et al. 2025) | (Kelly, et al. 2025) | (Blanc, et al. 2010) | (Cecchin, et al. 2019) | (Arriola, et al. 2018) | (Hamada, et al. 2018) |
| Retrieval | PRJEB104497 | GCA_002245815.2 | GCA_029339195.1 | GCA_047372815.1 | GCA_047372805.1 | GCA_000147415.1 | GCA_023343905.1 | GCA_002245835.2 | GCA_003063905.1 |
| Assembly length reported in source publication | 51.9* | 60.8 | 68.3 | 67.1 | 50.3 | 46.2 | 40.2 | 59.4 | 40.9 |
| Contigs | 30 | 300 | 1,499 | 20 | 227 | 414 | 44 | 159 | 82 |
| Largest Contig | 8,582,286 | 5,958,903 | 577,014 | 8,844,789 | 1,261,288 | 3,119,887 | 5,422,624 | 4,192,086 | 4,003,385 |
| Total Length | 62,572,437 | 61,018,900 | 68,293,531 | 67,114,787 | 50,300,000 | 46,159,512 | 40,437,440 | 59,566,223 | 40,934,037 |
| N50 (L50) | 3,831,832 (6) | 1,210,495 (13) | 115,803 (176) | 4,697,010 (6) | 3,623,000 (46) | 1,469,906 (12) | 2,825,136 | 2,592,956 | 1,727,419 |
| GC % | 67.36 | 67.2 | 65.17 | 65.22 | 65 | 67.14 | 61.6 | 63.95 | 70.46 |
| Ns per 100 kbp | 0 | 0 | 17.81 | 0 | 0 | 8,546.68 | 1,570.19 | 0 | 3,392.43 |
| Technology | PacBio, ONT & Illumina | PacBio & Illumina | HiSeq | PacBio | PacBio | Sanger | PacBio & Illumina | PacBio & Illumina | MiSeq & HiSeq |
| Coverage | 60-160x | 57x | 418x | 130x | 60x | 8.9x | 150x | 56x | 211x |
| Number of contigs with 2, 1, and 0 Telomeres | 9, 14, 8 | 4, 37, 259 | 2, 17, 1480 | 12, 4, 4 | 2, 54, 171 | 1, 17, 396 | 1, 11, 33 | 2, 21, 135 | 2, 11, 69 |
| BUSCO ODB10: |  |  |  |  |  |  |  |  |  |
| Eukaryota | C:92.5%[S:90.2%, D:2.4%],F:3.5%,M:3.9%,n:255 | C:88.6%[S:87.8%, D:0.8%],F:5.1%,M:6.3%,n:255 | C:89.4%[S:89.0%, D:0.4%],F:6.3%,M:4.3%,n:255 | C:93.3%[S:92.2%, D:1.2%],F:3.9%,M:2.7%,n:255 | C:90.2%[S:89.4%, D:0.8%],F:3.9%,M:5.9%,n:255 | C:78.8%[S:78.0%, D:0.8%],F:7.5%,M:13.7%,n:255 | C:93.3%[S:92.5%, D:0.8%],F:2.4%,M:4.3%,n:255 | C:90.6%[S:90.2%, D:0.4%],F:3.1%,M:6.3%,n:255 | C:80.0%[S:78.4%, D:1.6%],F:9.4%,M:10.6%,n:255 |
| Viridiplantae | C:93.2%[S:88.7%, D:4.5%],F:2.8%,M:4.0%,n:425 | C:90.1%[S:88.0%, D:2.1%],F:3.8%,M:6.1%,n:425 | C:91.3%[S:89.4%, D:1.9%],F:4.5%,M:4.2%,n:425 | C:94.6%[S:92.0%, D:2.6%],F:2.1%,M:3.3%,n:425 | C:91.3%[S:90.4%, D:0.9%],F:3.5%,M:5.2%,n:425 | C:84.5%[S:83.3%, D:1.2%],F:8.2%,M:7.3%,n:425 | C:93.9%[S:92.7%, D:1.2%],F:2.1%,M:4.0%,n:425 | C:92.7%[S:91.3%, D:1.4%],F:2.8%,M:4.5%,n:425 | C:78.6%[S:77.6%, D:0.9%],F:9.2%,M:12.2%,n:425 |
| Chlorophyta | C:99.1%[S:95.5%, D:3.6%],F:0.1%,M:0.9%,n:1519 | C:96.0%[S:94.3%, D:1.7%],F:0.7%,M:3.3%,n:1519 | C:98.5%[S:97.3%, D:1.2%],F:0.9%,M:0.6%,n:1519 | C:99.1%[S:97.0%, D:2.1%],F:0.2%,M:0.7%,n:1519 | C:97.1%[S:96.4%, D:0.7%],F:0.9%,M:2.0%,n:1519 | C:92.4%[S:91.2%, D:1.1%],F:3.1%,M:4.5%,n:1519 | C:99.0%[S:98.2%, D:0.9%],F:0.3%,M:0.7%,n:1519 | C:97.5%[S:96.6%, D:0.9%],F:0.6%,M:1.9%,n:1519 | C:90.1%[S:88.9%, D:1.2%],F:2.4%,M:7.5%,n:1519 |
| OMARK | Single: 1452<br>(80.62%) | Single: 1302<br>(72.29%) | Single: 1406<br>(78.07%) | Single: 1510<br>(83.84%) | Single: 1504<br>(83.51%) | Single: 1425<br>(79.12%) | Single: 1545<br>(85.79%) | Single: 1340<br>(74.40%) | Single: 1129<br>(62.69%) |
|  | Duplicated: 79<br>(4.39%) | Duplicated: 53<br>(2.94%) | Duplicated: 46<br>(2.55%) | Duplicated: 70<br>(3.89%) | Duplicated: 36<br>(2.00%) | Duplicated: 70<br>(3.89%) | Duplicated: 52<br>(2.89%) | Duplicated: 31<br>(1.72%) | Duplicated: 15<br>(0.83%) |
|  | Missing: 270<br>(14.99%) | Missing: 446<br>(24.76%) | Missing: 349<br>(19.38%) | Missing: 221<br>(12.27%) | Missing: 261<br>(14.49%) | Missing: 306<br>(16.99%) | Missing: 204<br>(11.33%) | Missing: 430<br>(23.88%) | Missing: 657<br>(36.48%) |
| PSAURON | 97.7 | 98.2 | 97.4 | 95.4 | 95.7 | 97.8 | 97.2 | 98.8 | 98.5 |
| Gene No. | 12,545 | 9,217 | 14,688 | 14,086 | 11,539 | 9,780 | 10,783 | 9,526 | 8,298 |
| Exon No. | 166,602 | 143,548 | 125,316 | 126,078 | 108,014 | 81,217 | 91,545 | 157,964 | 84,190 |
| Single Exon Genes | 167 | 68 | 817 | 576 | 394 | 233 | 562 | 65 | 191 |
| Intron No. | 151,083 | 128,472 | 110,648 | 111,992 | 96,475 | 70,995 | 80,376 | 142,318 | 74,610 |
| Mean Gene Size<br>(bp) | 3,696 | 5,247 | 3,671 | 3,410 | 3,408 | 2,927 | 3048 | 5,227 | 3,225 |

\*Estimated genome Size using LVgs (Mbp)

## Results and Discussion

### Genome Assembly and Annotation

Using a hybrid assembly approach described in the methods section, we generated a *M. conductrix* 186b genome assembly of 62.5 Mbp across only 30 contigs. To independently evaluate genome completeness, we used LVgs (Xing, et al. 2025) bioinformatic inference approach, resulting in an estimation of 51.9 Mbp. The mean coverage across the contigs using three different sequencing technologies is: 60X using Illumina reads (stdev 19.2); 168X using Pacbio HiFi reads (stdev 62.9); and 59X using Oxford Nanopore Technologies (ONT) reads (stdev 23.6). This assembly contained eight scaffolds of short length (<100 Kbp) that were not of organellar provenance. Five of these did not possess identifiable genes while three possessed low mapping coverage which varied by over three standard deviations from the mean coverage across the total assembly (i.e., Illumina coverage of 25.2X [stdev 11.1]; Pacbio HiFi coverage of 15.02X [stdev 19.15]; and Oxford Nanopore Technologies (ONT) coverage of 4.2X [stdev 2.22] – see Table S1). Collectively these eight contigs make up only ∼315 Kbp of the final assembly. The low sequencing coverage of the three short contigs suggest that the genome sequence sampled contains a minimal level of DNA contamination but potentially some accessory DNA elements. Accessory DNA elements often have variant copy number (e.g., ploidy counts) which may explain the variant coverage seen here when compared to the other 22 contigs sampled. This difference is then reflected in variant coverage scores. The inclusion of these variant contigs may lead to variant bioinformatic completion estimates, as seen in our LVgs analyses compared to the total assembly. The total genome contigs possessed a GC proportion of 67.36%. The assembly size and the GC% metrics are consistent with assemblies of other Chlorellaceae species (Table 1).

Next, we identified the number of candidate telomere regions incorporated within the assembly. This process identified nine contigs possessing a telomere at each end, suggesting these contigs encompass complete chromosomes. The assembly contained a further 14 contigs with only a single putative telomere at one end and then eight contigs without identifiable telomere-like repeat regions. These assembly data therefore suggest that *M. conductrix* 186b possesses sixteen chromosomes ([14/2]+9 = 16).

BUSCO (Tegenfeldt, et al. 2025) analyses of single copy genes demonstrated that the *M. conductrix* 186b assembly ranged from 92.5% to 99.1% completion. In contrast, OMARK (Nevers, et al. 2024) analyses identified a score of 80.2% completion (Table 1). These data combined with the telomere mapping data suggest that the *M. conductrix* 186b assembly shows a relatively high level of completion compared to the other algal genomes analysed (Table 1).

To explore how *M. conductrix* 186b relates to the eight published Chlorellaceae genomes compared here, we used FastANI (Jain, et al. 2018) to compute whole-genome average nucleotide identity using an alignment free method. These analyses demonstrated that *M. conductrix* 186b shares 99.87% identity with *M. conductrix* SAG 241.80, a green algal endosymbiont also isolated from a *Paramecium bursaria* strain. Similarly, mapping our Pacbio HiFi reads to the SAG 241.80 assembly identified a 99.7% alignment identity. FastANI comparisons to the other seven Chlorellaceae genomes identified between 76.45% and 77.70% alignment identity. Table S2 shows a pairwise alignment identity for all the genomes compared.

### Gene predictions

Gene predictions were carried out with the analyses pipeline BRAKER (Stanke, et al. 2006; Stanke, et al. 2008; Lomsadze, et al. 2014; Hoff, et al. 2016) using the RNA-seq and Iso-Seq transcriptome datasets independently. This process identified 12,545 genes using the RNA-seq data and 11,628 genes using the Iso-Seq data. Analysis of gene fusion profiles, discussed below, suggested that the RNA-Seq data identified an improved set of gene predictions when compared to the Iso-Seq analyses, which had an elevated recovery of longer genes potentially representing false fusions of ORFs. However, this disparity could be a product of a failure to accurately model alternative splicing events sampled in the long-read Iso-Seq transcript data and which are under-sampled in the short-read RNA-Seq methods.

The final set of 12,545 predicted genes included 134,839 exons (x̅ 10.7 per gene) and 122,306 introns (9.7 x̅ per gene), with a x̅ gene length of 3,688 bp (x̅ exon length of 140 bp, x̅ intron length of 224 bp). This set of gene predictions was used for gene annotation (described in the methods section below). Mapping a set of filtered and clustered transcripts (chimeras removed) for the combined dark and light RNA-Seq data to the CDS nucleotides of the gene predictions recovered a 96% hit rate using standard pBLAT setting (Wang and Kong 2019). This result is again consistent with the recovery of a near complete genome assembly.

### Identification of Organellar Genomes

MitoHiFi (Uliano-Silva, et al. 2023) was used to assemble and annotate the mitochondrial genome. The filtered dataset from the initial assembly was assembled using HiFiASM as part of the MitoHiFi pipeline along with annotations from KY629619.1 (a previously sequenced and publicly released *M. conductrix* mitochondrial genome of 74,708 bp which is predicted to encode 62 genes) (Fan, et al. 2017). This produced a putatively complete and circular mitochondrion of 75,292 bp (29.44% GC) featuring 60 predicted open reading frames; of which 27 tRNAs are present along with ATP6, & 8, COX I, II, & III, CYTB, NDP 1, 2, 3, 4, 5 & 6, atp 1, 4, & 9, nad 7, & 9, rpl 5, 6, & 16, and rps 2, 3, 4, 7, 10, 11, 12, 13, 14, & 19.

Similarly for the chloroplast, the program MitoHiFi was used in an experimental mode designed for chloroplast genome annotations using the *Micractinium simplicissimum* chloroplast genome (GenBank accession: NC_071969.1). This chloroplast genome possesses 114 genes across a 123,552 bp genome. This produced a partial chloroplast genome of 56,167 bp featuring 67 predicted open reading frames; of which 21 tRNAs are present along with; atp A, F, H & I, cemA, chlI, clpP, infA, pet A, B, D, G,& L, psaJ, psb B, C, D, H, I, M, N, & T, rpl 2, 5, 12, 14, 16, 19, 20, 23, & 36, rpo A, B, & C1, rps 2, 3, 7, 8, 9, 11, 12, 18, & 19, 6ufa, ycf 3, & 4.

### Investigating patterns of gene fusion prediction across multiple Chlorellaceae genome assemblies

To select a set of *M. conductrix* 186b gene predictions to retain for further use, we initially compared our RNA-seq and Iso-Seq informed results with the publicly available gene repertoire for *M. conductrix* SAG 241.80 (REF). However, despite ∼99.9% nucleotide identity between the two genome assemblies, this revealed large disparity in predicted gene content, with both sets of *M. conductrix* 186b gene predictions containing many more genes than predicted for *M. conductrix* SAG 241.80 (*M. conductrix* 186b RNA-Seq informed gene predictions: 12,545; *M. conductrix* 186b Iso-Seq informed gene predictions: 11, 628; *M. conductrix* SAG 241.80: 9,217). Moreover, mean gene length was notably higher for *M. conductrix* SAG 241.80 (*M. conductrix* 186b RNA-Seq informed gene predictions: 3,696 bp; *M. conductrix* 186b Iso-Seq informed gene predictions: 3,919 bp; *M. conductrix* SAG 241.80: 5,247 bp), indicating frequent discrepancies between studies in whether sequences were assigned to single or multiple ORFs.

We explored this further by generating and comparing gene fusion profiles for the three sets of *M. conductrix* gene predictions and seven additional Chlorellaceae datasets (Table 1) using the bioinformatic pipeline ‘fusion duplication finder BLAST’ (fdfBLAST) (Leonard and Richards 2012). Briefly, fdfBLAST runs a set of all-against-all gene repertoire-to-repertoire comparisons using reciprocal best BLAST hits. The pipeline then applies a custom algorithm to detect multiple genes from one genome that map to a single ORF in an alternative genome where the genes have minimal spatial overlap within the ‘fused gene’ ORF. Recovered ‘hits’ could represent true gene fusions or genome annotation artefacts produced by erroneously amalgamated gene predictions. The pipeline assigns a score to the asymmetric hits identified based on overlap in the ORF maps, enabling users to profile and sort candidate differentially distributed gene fusions for further analyses.

Estimated rates of candidate gene fusions ranged from 5.6 – 21.5% across the 10 Chlorellaceae gene repertoires compared, with the lowest and highest rates obtained for the *M. conductrix* 186b RNA-Seq informed predictions and *M. conductrix* SAG 241.80, respectively (Fig. 1B). Whilst some may reflect true fusions, many of the gene fusions differentially distributed across the Chlorellaceae gene repertoires are likely to represent artifacts, i.e., products of gene calling methodologies not optimized for non-model organisms such as Chlorellaceae in combination with the historical use of patchy transcriptome sequencing datasets which, for example, poorly account for alternative splicing. Indeed, in the analysis of *M. conductrix* 186b, detected rates of putative gene fusions varied from 5.6 to 9.8% depending on the type of mRNA sequencing conducted (RNAseq vs. Iso-Seq), indicating a bias towards recovery of fused ORFs in the Iso-Seq data. We selected to retain the RNA-seq informed *M. conductrix* 186b gene predictions for further use, reasoning that the lower rate of differentially distributed gene fusions made this the more parsimonious option in terms of false gene fusion prediction.

**Figure 1.**
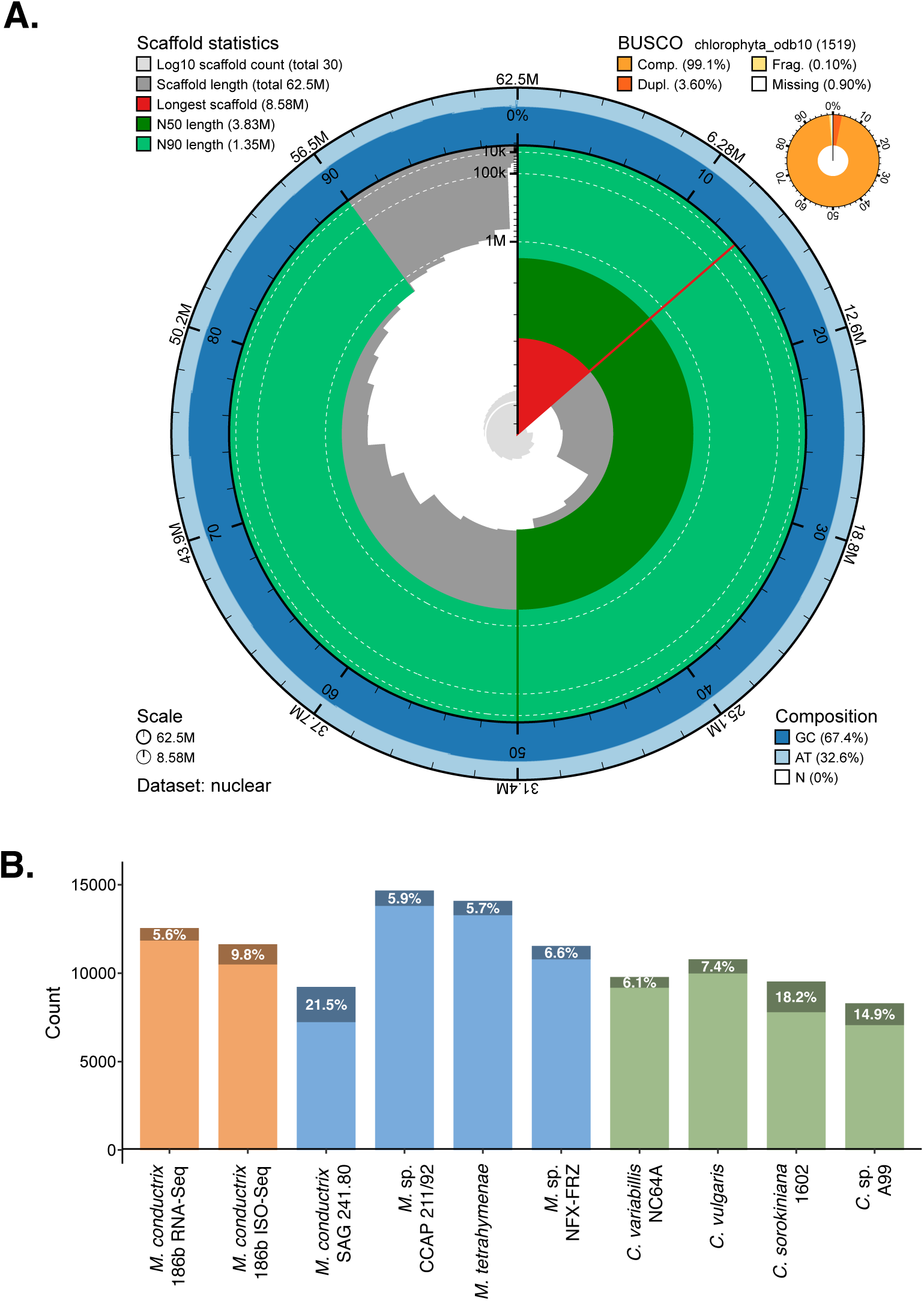
**A.** A snail plot generated by BlobToolKit (Challis, et al. 2020) showing the general statistics for the genome assembly. Total scaffold length in grey, with N50 and N90 in shades of green. Also reported are the BUSCO chlorophyta_odb10 results (orange circle), and genome composition as GC% **B** Bar plots showing the number of predicted genes and the percentage of putative gene fusions across nine taxa (*Micractinium* species in blue, *Chlorella* species in green), along with the two sets of gene predictions generated for the 186b strain (in orange) using the RNA-seq or ISO-seq data.

## Conclusion

Here, we report a highly complete genome assembly for the endosymbiotic alga *M*. *conductrix* 186b. This complements our recent publication of the *P. bursaria* 186b host genome and will facilitate further studies focussing on mechanisms underpinning endosymbiotic interactions, such as comparative genomics analyses and laboratory evolution or genetic modification experiments. Our analyses demonstrate high variability in putative gene fusions across predicted Chlorellaceae gene repertoires, highlighting the necessity of checking ORF predictions in downstream studies of target genes within this group.

## Data Availability

All sequencing reads are deposited at EMBL EBI ENA with the BioProject ID PRJEB104497. Genomic PacBio Reads ERR16000047, Genomic ONT Reads ERR16000086:ERR16000092 and Genomic Illumina Reads: ERR15999986. Transcriptomic Reads: ERR16000280:ERR16000285, PacBio Iso-Seq Reads ERR16000279. The Mc 186b genomic assembly is available at CDWRBR01. The complete mitochondrion sequence can be accessed at ERZ28769978, and the partial chloroplast genome can be accessed at ERZ28769979. The assembled genome sequences and annotations can also be accessed from Github and/or Zenodo. An update version of fdfBLAST adapted for use with DIAMOND is available at https://github.com/guyleonard/fdfBLAST/releases/tag/1.2.

## Funding

This work was funded by grants from NERC (NE/V000128/1) to MAB and DDC and the BBSRC (BB/X016439/1) to MAB, DDC and TAR. TAR was supported by a Royal Society University Research Fellowship (UF130382) and latterly by an ERC Consolidator Grant (CELL-in-CELL).

## Acknowledgements

We acknowledge David Hopkins for contributions to the lab work and staff at the NERC Environmental Omics Facility at University of Sheffield for their assistance with method development.

## Materials & Methods

### DNA Extraction and Sequencing

Symbiotic algae were isolated by washing 186b *Paramecium* cells with NCL with ampicillin, lysing cells by sonication (20% power, 8 s), and plating the lysate on either Modified Bold Basal Media (MBBM, Culture Collection of Algae and Protozoa [CCAP] Available at: https://www.ccap.ac.uk/wp-content/uploads/MR_MBBM.pdf [Accessed 25 May, 2021]), or modified artificial WC media (Lebret, et al. 2012) supplemented with amino acids, each with 1.5 % agar and ampicillin. Plates were incubated at 20°C with 33 µE m⁻² s⁻¹ of light for longer term algal maintenance and at 14:10 light: dark cycle, for growing biomass and at 25°C with 50µE m⁻² s⁻¹ of light at a 14:10 light: dark cycle for growing biomass. After 3—4 weeks, individual colonies were moved to standard algal growth conditions in liquid MBBM (25 °C, 50 µE m⁻² s⁻¹ light, 14:10 L:D cycle, 110 rpm shaking).

Strain identity was confirmed by sequencing SSU rRNA and ITS2 encoding gene regions (Hoshina, et al. 2005). Cultures of the 186b algal isolate were maintained at 20 °C under 33 µE m⁻² s⁻¹ of light with a 14:10 light:dark cycle. Prior to DNA extraction, 5 mL of culture was transferred to 25 mL of fresh MBBM medium and grown for 14 days. The culture was then centrifuged at 30,000 × g for 10 minutes in a 50 mL Falcon tube and flash-frozen in liquid nitrogen.

For PacBio long-read sequencing, 60 mg of the flash frozen sample was ground in a pre-chilled mortar and pestle for 10 minutes, with liquid nitrogen added as needed to maintain freezing. DNA was then extracted using the NucleoBond HMW DNA extraction kit (Macherey Nagel) and eluted in 200 µL of HE buffer. The extracted DNA was shipped on ice to the Centre for Genomic Research (CGR) at the University of Liverpool, where it was cleaned using AMPure XP beads and sequenced using PacBio HiFi technology on the Sequel II SMRT Cell platform in CCS run mode.

For short-read sequencing, a custom CTAB-ethanol precipitation protocol was used. Samples were homogenized with 0.5 mm zirconium oxide beads in 900 µL of CTAB solution, followed by incubation with 50 µL of proteinase K at 65 °C for 30 minutes. Subsequently, 3 µL of RNase A was added, and the mixture was incubated at 37 °C for 30 minutes. A 25:1 chloroform:isoamyl alcohol solution was added, samples were inverted 10 times, and centrifuged to isolate the aqueous phase. DNA was pelleted by centrifugation at 10,000 rpm for 60 minutes at 4 °C, washed twice with 70% ice-cold ethanol, and re-eluted in 200 µL of TE buffer. Samples were sent to CGR at the University of Liverpool for short-read sequencing using the Illumina NovaSeq 6000 platform, using TruSeq PCR-free kit using a single lane of a S4 flow cell.

For ONT sequencing DNA was isolated using the same methods as for the PacBio analyses. Libraries were prepared using the Ligation Sequencing Kit SQK-LSK114 (Kit 114 chemistry) following the manufacturer’s protocol. Libraries were run using FLO-MIN114 (R10.4.1) flow cells on a GridION X5 Mk1 (GXB02152 – with the following settings: run length of 72 hrs, active channel selection on, pore scan frequency of 1.5 hrs, reserved pores on, read splitting on, minimum read length of 1000 bp and base-calling in the high-accuracy mode.

### RNA Extraction and Sequencing

*M. conductrix* 186b cultures were grown to mid-late log phase in MBBM at 24°C with a 14:10 light:dark cycle then were processed as below after six hours of light exposure or six hours after covering with foil to prolong the dark cycle. Cultures were centrifuged at 2800 × g for 5 minutes at 4°C, flash-frozen in liquid nitrogen and then maintained at -80°C until further processing. Pellets were briefly thawed on ice then were resuspended in 800 µl Monarch® DNA/RNA protection reagent (New England Biolabs) and transferred to BeadBug^TM^ tubes prefilled with 0.1 mm TriplePure zirconium beads (Merck). Cells were homogenised with a TissueLyser II (Quiagen) using three consecutive 90 sec runs at top speed (30 Hz), with incubations on ice for 1 min between each run, then were placed back at -80°C until further processing. Total RNA was isolated using a Monarch Total RNA Miniprep Kit following the manufacturer’s protocol. RNA quantity and integrity were assessed on an Agilent 2100 Bioanalyser using an Agilent RNA 6000 Nano kit following the manufacturer’s protocol. Purity was assessed using a NanoDrop^TM^ spectrophotometer (Thermo Scientific). Iso-Seq sequencing was performed by Genewiz for two RNA samples with sufficiently high RNA integrity scores (both from cultures exposed to light) using a single SMRT cell on a PacBio Revio Long-Read Sequencer. RNA-seq sequencing was performed by Genewiz for six RNA samples (three light, three dark) with PolyA selection using the Illumina® NovaSeqTM X series platform.

### Iso-Seq Assembly

The PacBio Iso-Seq library was generated from two RNA samples from two culture replicates sampled during the light phase. These datasets were analysed using the standard isoseq.how v4.0 CLI Workflow. Briefly, adapters and barcodes were removed with “lima”, which is followed by a refining process and a clustering algorithm (isoseq refine and isoseq cluster2, respectively). This converts the raw data (23,991,521 reads and 21,776,230 reads) into full-length non-concatemer transcripts (11,294 transcripts comprising 29.28 Mbp). To assess the coverage of the Iso-Seq transcriptome, we used pBLAT (Wang and Kong 2019) to search the transcript CDS (10,065) against the genome, resulting in 100% coverage.

### RNA-Seq Assembly

Six libraries (three light, three dark) were assembled together using Semblans (Woodcock-Girard, et al. 2025) – a pipeline including read correction, QC and trimming, assembly using Trinity, chimera detection, clustering and gene prediction using TransDecoder all in one. This produced 82,448 transcripts producing 46,890 CDS sequences. These mapped at a rate of 9X to the RNA based gene predictions (see below).

### Genome Assembly

The FASTQ files were recovered from CGR Liverpool and an initial assembly using the PacBio HiFi reads was created with HiFiASM (Cheng, et al. 2021). This produced an assembly with 1,085 contigs, a total length of 101,151,861 bp and an N50 of 1,786,873 bp. Subsequently, this assembly was analysed using BLOBTOOLKIT (see Fig. S1 & (Kumar, et al. 2013)). This showed three separate sets or ‘blobs’: a smaller set of large blobs with a relatively consistent coverage range with a GC content of around 65%; along with two larger sets of much smaller blobs with ∼30% and ∼34% GC respectively. This looked highly suggestive of nuclear, mitochondrial and chloroplast content previously reported in (Arriola, et al. 2018) and from other Chlorophyta genomes sampled in Table 1. Given this distribution and association, the scaffolds were filtered accordingly (using the BLOBTOOLS filter command and the relevant GC cutoffs) into three subsets, the putative nuclear, mitochondrion and chloroplast genomes. Original HiFi reads were then extracted using SAMTOOLS for each data set.

The nuclear set was then re-assembled using HiFiASM and this time supplying the standard Chlorophyta telomeric repeat of CCCTAAA_n_, which produced an assembly with 30 contigs containing a total length of 62,572,437 bp, an N50 of 3,831,832 bp, a L50 of 6, and a GC content of 67.36 %. This is similar to other Chlorophyta assemblies (see Table 1), and supports the use of the separate ‘blob groups’ for distinct genome assemblies.

Coverage depth was calculated for all contigs and sequencing technologies (Illumina x̅ 60X - stdev 19.2, PacBio HiFi x̅ 168X - stdev 62.9, & ONT x̅ 59X -stdev 23.6) using MINIMAP2 (Li 2018, 2021) and SAMTOOLS (Danecek, et al. 2021) which identified three contigs with very low coverage compared to the other contigs (Table S1). These three contigs are retained but could potentially represent contamination or extra accessory genomic DNA sequences. Nonetheless, these contigs were kept in the published assembly and labelled as variant.

The genome assembly was polished using MEDAKA (https://github.com/nanoporetech/medaka) for one round, incorporating the Oxford Nanopore Technology reads, and one round of polishing using PILON (Walker, et al. 2014) incorporating the Illumina reads. Post assembly and polishing, the contigs were repeat masked using RepeatModeller and RepeatMasker which soft masked 21.5% of the genome as repeat sequences. 13.6% were identified as interspersed repeats (∼11.9% unclassified and ∼1.2% retroelements) while 8.26% were identified as simple repeats

Further to the analysis of repeat motifs, we used TAPESTRY (Davey, et al. 2021), quartet (Lin, et al. 2023), and BLAST+ (Camacho, et al. 2009) searches to identify regions of the assembly that represent telomeres. This identified nine contigs with telomere-like signal at both ends, thirteen at one end, and eight with no detectable signal. The program TELOMORE (initially developed for the analysis of yeast genomes, (https://github.com/dalofa/telomore) was utilised to try and extend telomere signals for the contigs recovered specifically using the ONT data. Whilst it was able to extend telomere signals partially on most contigs where they had previously been detected, it was only able to add a telomere signal to the end of one additional contig. This suggests a final putative chromosome number of 9 + (14 / 2) = 16.

Genome size estimation was carried out with LVgs (Kokot, et al. 2017; Ranallo-Benavidez, et al. 2020; Cheng, et al. 2024; Xing, et al. 2025), a k-mer based pipeline that utilises HiFi reads, and suggests a genome size of 51.9 Mbp using a k of 27 bp - an estimate of about 10 Mbp smaller than the total assembly size reported here (62.7 Mbp). BUSCO (Tegenfeldt, et al. 2025) and OMARK (Nevers, et al. 2025) analyses were completed with Eukaryota, Viridiplantae, Chlorophyta (all ODB10) and Chlorophyta (from the LUCA OMA database), respectively. Other genome statistics were carried out using QUAST (Gurevich, et al. 2013) (see Table 1) and BLOBTOOLKIT (Challis, et al. 2020) (see Fig. 1A).

### Gene Prediction and Annotation

Gene predictions were carried out with BRAKER (Stanke, et al. 2006; Stanke, et al. 2008) with the raw RNA-Seq sequencing data, BUSCO identified ‘chlorophyta’ gene-set, and the masked genomic assembly. A supplementary gene prediction set was also completed by extracting the intron positions from the assembled Iso-Seq transcripts and utilising a modified version of BRAKER that accepts long reads (Bruna, et al. 2024).

Testing the two sets of gene predictions with PSAURON (Sommer, et al. 2025) – which assigns a score to gene predictions based on a Machine Learning model – indicated a score of 96.8 for the Iso-Seq intron based BRAKER predictions and a score of 97.7 for the RNA-Seq based BRAKER. Genes were also tested for putatively fused ORFs using the software fdfBLAST (Leonard and Richards 2012). Manual checks of gene predictions demonstrated there were an over-abundance of candidate gene fusions (see fdfBLAST section below). This process also indicated the use of RNA-Seq data identified a lower count for putative gene fusions than the Iso-Seq approach (Fig. 1B).

Functional annotation was added to the gene predictions using multiple databases from InterProScan (Jones, et al. 2014) (PFAM, EGGNOG, and BUSCO) and integrated by the FUNANNOTATE (Palmer and Stajich 2020) pipeline using the “other” mode.

### Gene Fusion Detection

A modification of the program fdfBLAST (https://github.com/guyleonard/fdfBLAST/releases/tag/1.2) with the script updated to use DIAMOND BLAST (for computational speed (Buchfink, et al. 2021)) was used with the predicted proteins from the eight published algal genomes in Table 1 and the two sets of gene predictions produced from the 186b genome assembly as source data using either the RNA-Seq or the Iso-seq datasets.

## Figures and Figure Legends

**Supplementary Figure 1.**
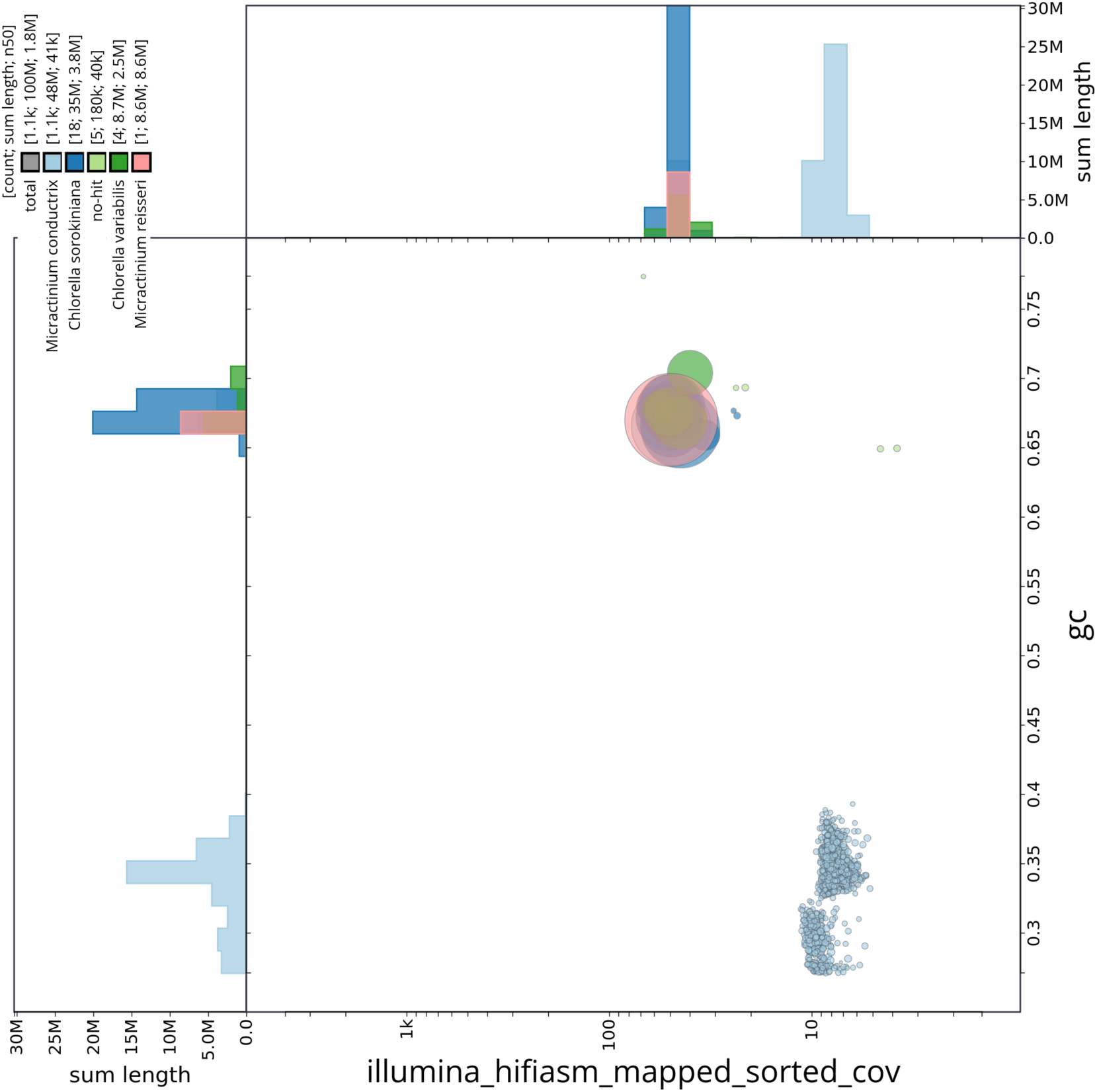
A Blobplot showing the distribution of contigs based on their GC content and coverage (from Illumina reads), blob sizes are indicative of contig size. Colours are assigned from blast taxonomy. It shows three distinct groups of blobs, the larger blobs around 0.65% GC are indicative of the nuclear genome, whereas the two other blobs are suggestive of the mitochondrial and chloroplast genomes.

**Supplementary Figure 2.**
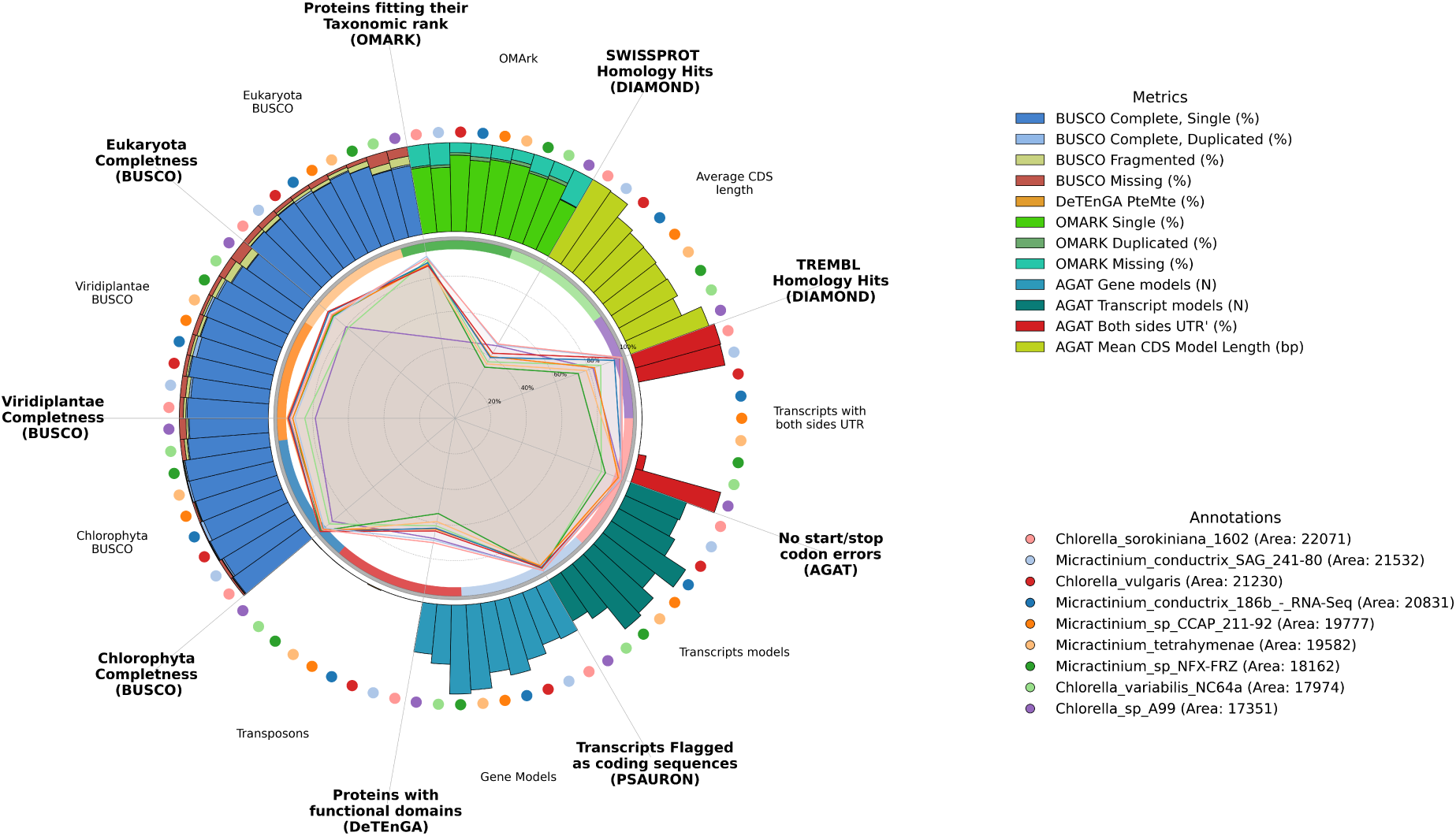
A GAQET plot showing various metrics for each of the nine taxa, including the *M. conductrix* 186b RNA-seq informed gene prediction set. Taxa are represented by colours in the key, with radial bar charts per analysis e.g., BUSCO completion, OMARK rank and homology hit rate to Swissprot/TREMBL.

**Supplementary Table 1.** This table holds information on the size and coverage data for each contig produced in the final nuclear assembly. It also shows number of genes predicted per contig and telomere locations.

**Supplementary Table 2.** Pairwise genome alignment percent identity across the nine Chlorellaceae genomes analysed.

## Notes

### Competing Interest Statement

The authors have declared no competing interest.

